# Empowering bioinformatics communities with Nextflow and nf-core

**DOI:** 10.1101/2024.05.10.592912

**Authors:** Björn E. Langer, Andreia Amaral, Marie-Odile Baudement, Franziska Bonath, Mathieu Charles, Praveen Krishna Chitneedi, Emily L. Clark, Paolo Di Tommaso, Sarah Djebali, Philip A. Ewels, Sonia Eynard, James A. Fellows Yates, Daniel Fischer, Evan W. Floden, Sylvain Foissac, Gisela Gabernet, Maxime U. Garcia, Gareth Gillard, Manu Kumar Gundappa, Cervin Guyomar, Christopher Hakkaart, Friederike Hanssen, Peter W. Harrison, Matthias Hörtenhuber, Cyril Kurylo, Christa Kühn, Sandrine Lagarrigue, Delphine Lallias, Daniel J. Macqueen, Edmund Miller, Júlia Mir-Pedrol, Gabriel Costa Monteiro Moreira, Sven Nahnsen, Harshil Patel, Alexander Peltzer, Frederique Pitel, Yuliaxis Ramayo-Caldas, Marcel da Câmara Ribeiro-Dantas, Dominique Rocha, Mazdak Salavati, Alexey Sokolov, Jose Espinosa-Carrasco, Cedric Notredame, the nf-core community.

## Abstract

Standardised analysis pipelines are an important part of FAIR bioinformatics research. Over the last decade, there has been a notable shift from point-and-click pipeline solutions such as Galaxy towards command-line solutions such as Nextflow and Snakemake. We report on recent developments in the nf-core and Nextflow frameworks that have led to widespread adoption across many scientific communities. We describe how adopting nf-core standards enables faster development, improved interoperability, and collaboration with the >8,000 members of the nf-core community. The recent development of Nextflow Domain-Specific Language 2 (DSL2) allows pipeline components to be shared and combined across projects. The nf-core community has harnessed this with a library of modules and subworkflows that can be integrated into any Nextflow pipeline, enabling research communities to progressively transition to nf-core best practices. We present a case study of nf-core adoption by six European research consortia, grouped under the EuroFAANG umbrella and dedicated to farmed animal genomics. We believe that the process outlined in this report can inspire many large consortia to seek harmonisation of their data analysis procedures.

## Introduction

Advancements in large-scale molecular biology methods have driven an unprecedented increase in data generation^1–3^. The volume of data generated pushes computational capacities to their limits, revealing an urgent need for robust and scalable data analyses^4^. Workflow management systems (WfMSs) are now the recommended solution when dealing with high-throughput data analysis pipelines^5^. While some frameworks such as the Galaxy framework^6^ focus on user-friendly graphical interfaces, tools such as Snakemake^7^ and Nextflow^8^ are designed for bioinformaticians familiar with programming. They combine the expressiveness^9^ of Bash with additional features to better support reproducibility, traceability, parallelisation, and portability across different infrastructures (HPC clusters, cloud computing, and workstations). When considering some of the most popular WfMSs^9^ we found that there were more than two and half times the number of citations in 2023 than in 2018 (Fig. 1, Supplementary Fig. 1, Supplementary Table 1, Supplementary Table 2). A notable trend over the last six years is a decrease in Galaxy citations in favour of Snakemake and Nextflow. In 2023, Nextflow was the resource with the highest usage growth and in that same year, Google Scholar data indicates that Galaxy and Nextflow each accounted for about a third of reported WfMS usage.

**FIGURE 1:**
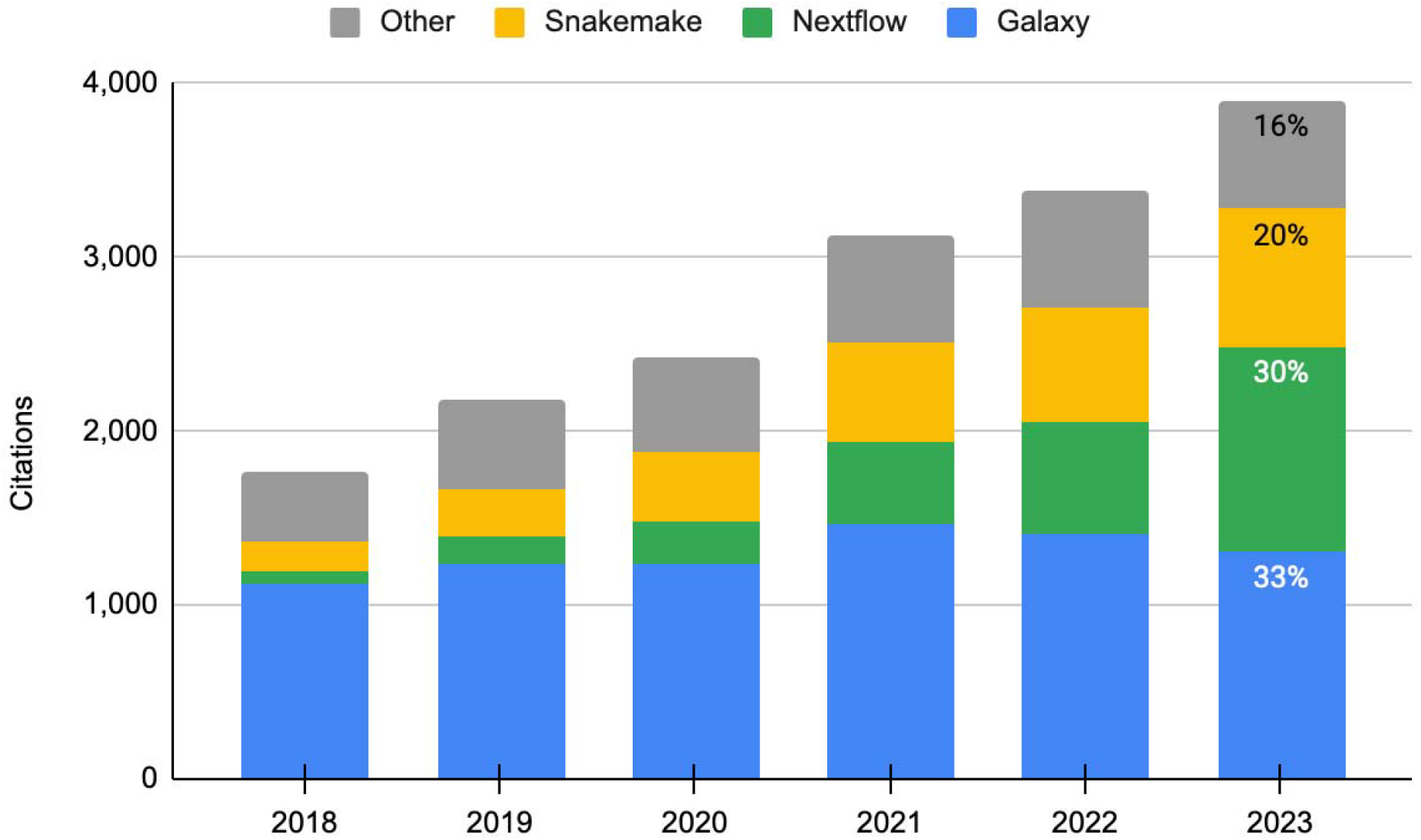
Google Scholar citation counts for bioinformatics workflow management systems. Sum of citations of the major publications of Galaxy, Nextflow, and Snakemake between 2018 and 2023 (Data in Supplementary Table 1).

While WfMSs provide a powerful way to bundle available methods into pipelines, they do not provide standards for how this bundling should be carried out. This gap has led to the establishment of pipeline registries for Snakemake (Snakemake Workflow Catalog (SWC), https://snakemake.github.io/snakemake-workflow-catalog/; Snakemake-Workflows, https://github.com/snakemake-workflows) and Nextflow (nf-core, https://nf-co.re/). These workflow repositories enforce implementation guidelines, effectively providing best-practice standards for both WfMSs. The nf-core community has been a key driver behind Nextflow adoption, in part due to the excellent quality of its curated pipeline collection. A recent study found that 83% of nf-core pipelines could be deployed and run automatically without a crash, nearly 4x better than other catalogues^10^.

### Nextflow and nf-core: recent developments

#### Expansion of the nf-core pipeline community

The nf-core community was created in 2018 as a curated collection of pipelines implemented according to agreed-upon best-practice standards^11^. These off-the-shelf pipelines are characterised by reproducibility, standardisation, and rapid result generation. There are now nearly 100 pipelines available within nf-core, covering the analysis of a broad range of data types, including high-throughput sequencing, mass spectrometry, protein structure prediction, microscopy, satellite imagery analysis, and economics. These curated pipelines are supported by over 2,000 contributors and developers on GitHub, over 100 institutions, and over 8,000 users on nf-core’s primary communication platform Slack. Pipeline long-term maintenance is achieved by an efficient cooperation model allowing teams across sites to collectively and simultaneously contribute according to their needs and capacity (Fig. 2a).

**FIGURE 2:**
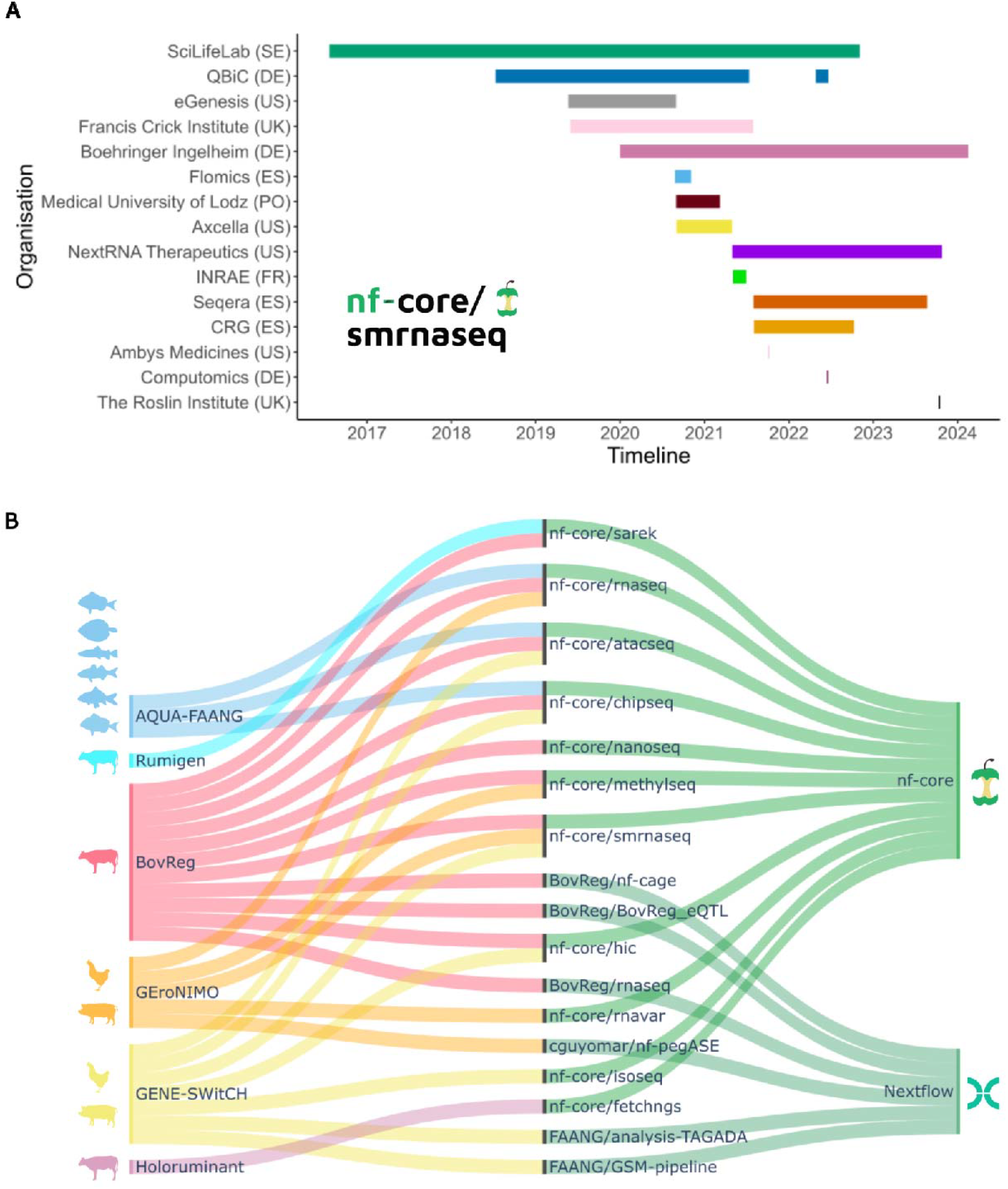
Pipeline maintenance and usage. (A) Major contributions to the *nf-core/smrnaseq* pipeline over time by different academic institutions or private companies. Data for individual contributors is collapsed to their institution (SciLifeLab: 3; QBiC: 2; Boehringer Ingelheim: 3; Seqera: 4; all the others: 1) (B) Nextflow analysis pipelines used in the EuroFAANG consortia for the functional annotation of various species’ genomes.

#### Broadened nf-core outreach

nf-core organises programs and events to facilitate sustainable growth of the community. Since its inception, nf-core has run 13 virtual, hybrid, or in-person hackathon events that have attracted global attendance. Nearly 100 ‘nf-core/bytesize’ webinars describing community advancements, coding approaches, and pipelines, have been shared online, attracting tens of thousands of views. A mentorship program pairs experienced Nextflow and nf-core developers with new community members from underrepresented groups. Free community Nextflow and nf-core training videos are recorded and shared online every 6 months, accounting for over 35,000 views on YouTube since October 2022 (Supplementary Table 4). nf-core strives for accessible community programs and events, irrespective of language, geography, time zones, and the ability to travel.

#### Evolution of the Nextflow workflow management system

The most notable development in Nextflow during the past five years has been the introduction of the DSL2 syntax. This introduced the ability to define complex workflows which can be organised into reusable components. This has dramatically improved the encapsulation and re-use of analytical steps shared across pipelines. Nextflow has also benefited from improved support for cloud-based computing platforms, support for on-demand container provisioning via the Wave service^12^, and extended support for many HPC schedulers and software container engines (15 schedulers and 7 container engines, as of version 23.10.0). Community-driven projects such as nf-test^13^, a testing framework for Nextflow pipelines and components, have garnered rapid adoption. Notably, Nextflow plugins ease the extension of core Nextflow functionality, illustrative examples are nf-validation^14^ (JSON schema validation of pipeline parameters and sample sheets), nf-co2footprint^15^ (energy consumption and carbon intensity from pipeline compute), and nf-prov^16^ (provenance reports for pipeline runs). This wide range of improvements may explain why Nextflow is being adopted in fields beyond biological applications, with new pipelines in fields as diverse as astrophysics^17^, earth science^18^, and economics^19^.

#### Impact of module support

The main impact of Nextflow’s DSL2 transition on nf-core has been to enable the establishment of a shared repository for standardised modules and subworkflows. The *nf-core/modules* repository currently features more than 1000 modules and 50 subworkflows, which are used by nf-core and non-nf-core pipelines. Each module is a wrapper around a single tool function and bundles the required software via Docker / Singularity images and a Conda environment, provided via the Bioconda^20^ and Biocontainers communities^21^. In this way, modules encapsulate the correct environment for each embedded software, thus mitigating dependency conflicts between software packages. nf-core also supports self-hosted module repositories. This allows direct integration of internal development with community-maintained modules and subworkflows, thereby bridging closed- and open-source development.

#### Improved pipeline standardisation

Adopting a unified pipeline structure and incorporating shared features guarantees consistent functionality and high-quality standards across nf-core pipelines. For this, nf-core has developed a base template for pipelines. The template receives regular updates which are semi-automatically synchronised across all pipelines. The template establishes pipeline file structure, with code, documentation, and continuous integration (CI) tests. Pipelines are tested using automated linting tests for code formatting and syntax, and flags outdated nf-core modules. To guarantee deployability, each pipeline comes with minimal datasets used to test the pipeline for every pull request. CI testing also includes a “full-size” dataset with a realistic input to test portability to diverse platforms for each stable release, executed automatically on AWS and Microsoft Azure after every pipeline release. The output files are publicly accessible and can be browsed on the nf-core website.

#### Extended tooling

The *nf-core/tools* package provides helper tools to improve accessibility for users of any Nextflow pipeline. Functionality includes commands to download pipelines and associated software for offline use and methods for launching pipelines, including a command-line and a graphical web-browser-based pipeline-launch interface to enable users less comfortable with command-line interfaces. The CLI is a key part of nf-core development, with functionality to create new pipelines based on the nf-core template, create and modify JSON schema files for parameter validation/documentation. Subcommands allow developers to discover, install, and update modules and subworkflows, as well as create new ones from a template. Automation synchronises existing pipelines with new template versions, as well as updating modules and subworkflows to the latest version of the tool. A lint command tests pipelines and components for adherence to nf-core guidelines.

#### Reproducibility and data provenance

Achieving full reproducibility when running a pipeline is an essential requirement of the FAIR paradigm. Nextflow and nf-core simplify this task by supporting the programmatic collection of execution parameters (eg. tool versions, command-line flags) through parameter files, the nf-prov plugin and MultiQC^22^ summary reports. Provenance is especially critical when comparing results across large consortiums, such as EuroFAANG (see next section). These features were essential to meet the FAANG metadata standards^23^, facilitating comparability, reproducibility, and standardisation.

### Uptake of nf-core by the farmed animals genomics community

#### Functional Annotation of Animal Genomes project (FAANG)

FAANG^24^ (Functional Annotation of Animal Genomes) is a coordinated international effort established in 2014, with the explicit purpose of harmonising genotype-to-phenotype research in farmed animals. EuroFAANG is a formalised initiative of FAANG partnering organisations across Europe, which currently encompasses six of EU’s Horizon 2020 program consortia: AQUA-FAANG, BovReg, GENE-SWitCH, RUMIGEN, GEroNIMO, and HoloRuminant. These six consortia bring together more than 80 academic institutions and commercial companies spread across Europe, North America, Middle East, and Australia. While each project specialises in a distinct set of farmed animal species (multiple fish species, cattle, pig, and chicken), they share the common goal of improving our understanding of the genome-phenome relationship through the comprehensive annotation of genomic functional features. This collective effort systematically captures dynamic gene expression, chromatin accessibility, and epigenetic state under a range of biological conditions. Comprehensive functional annotation demands the generation and analysis of large sequencing datasets derived from distinct functional assays including RNA-seq, ATAC-seq, and ChIP-seq. Together, these projects are committed to implementing an analysis strategy adhering to the FAIR (findability, accessibility, interoperability, and reusability) principles of scientific reproducibility^25,26^.

The projects AQUA-FAANG, BovReg, and GENE-SWitCH were the first to use the EuroFAANG name as a common umbrella. EuroFAANG is now developing as a future European Research Infrastructure^27^ to bring together national facilities at the pan-European level, in the fields of animal genetic resources, phenotyping and breeding, and animal health. The EuroFAANG Research Infrastructure project builds on the foundation provided by the six current H2020 EuroFAANG projects and connects with existing infrastructures for data management and animal agriculture and aquaculture in the European research infrastructure landscape. It also addresses the research priorities outlined in FAANG to Fork^28^, which mirrors the European Commission’s European Green Deal Farm to Fork strategy. The recommendation, adoption, adaptation, and reuse of nf-core workflows for future genotype-to-phenotype research remains a key aspect of the goals of the EuroFAANG Research Infrastructure, to ensure that analyses conducted across European institutes and industry are interoperable, standardised, and reusable.

#### Necessity of interoperability and standardisation

The use of nf-core provides an effective standardisation strategy, which avoids technical differences that can be introduced by using varied parameterisation and analysis tools. Such standardisation ensures the comparability of results. For instance, AQUA-FAANG relied on nf-core pipelines to exploit the evolutionary conservation of genomic regulatory elements and epigenetic states across six farmed fish species for multiple biological conditions and matched tissue panels^29^. Data integration across projects is critical, as projects such as the farm animal GTEx^30–32^ are expected to keep integrating a larger number of species and deeper sequencing over time. Interoperability, achieved through the use of a common analytic framework is therefore an essential component of long-term sustainability, well beyond the goal and lifetime of projects like EuroFAANG.

#### Main hurdles towards a common bioinformatics analysis framework

Establishing common standards is still one of the most challenging goals of bioinformatics^33^. Research groups often have established methods and adopting new standards may require re-training staff, as well as re-implementing and re-validating methods already in production. The process of standards adoption can benefit from being spear-headed by a designated leader. When the AQUA-FAANG consortium decided to utilise nf-core, the University of Edinburgh and the Norwegian University of Life Sciences took the responsibility of deploying the framework and training the other AQUA-FAANG groups to use nf-core pipelines. Regardless of which framework is chosen, the transition roadmap is as important as the endpoint – a gradual transition is more likely to succeed in avoiding fragmentation across groups.

#### Choosing a bioinformatics analysis framework

Discussions and surveys about methods, languages, and standards were carried out among the EuroFAANG partners (Table 1). These consultations reflected the predominance of Nextflow as a favoured WfMS, and identified the needs of consortia that were not already met by the methods available in nf-core. We found that the 15 consortium partners involved in data analysis were carrying out 78 analyses, using four different programming languages: Nextflow (70), Snakemake (3), Bash (4), and Perl (1). A key feature of Nextflow is its lack of language dependency. Even though the Nextflow DSL is based on Groovy, the scripts and functions of a pipeline can be directly ported into Nextflow in their original language. This enables pipelines to be gradually ported into Nextflow without the need to rewrite scripts or do extensive revalidation. For example, the BovReg eQTL analysis workflow was initially written in Bash and was ported into a Nextflow pipeline. The wide range of analyses available in nf-core and its fast-growing community are important aspects of long-term pipeline maintenance.

#### EuroFAANG pipeline development within the nf-core ecosystem

All nf-core pipelines are open source and welcome new contributions. For instance, EuroFAANG partners have been heavily involved in the maintenance and development of several existing nf-core pipelines, including *nf-core/atacseq, nf-core/chipseq*, and *nf-core/smrnaseq* (Table1). PacBio-based genome annotation was identified as a gap for the GENE-SWitCH project, leading one of its partners (Roslin Institute, University of Edinburgh, UK) to port their Iso-Seq Nextflow pipeline into nf-core (*nf-core/isoseq*^34^). Similar collaborative efforts have been initiated within GEroNIMO and BovReg for eQTL analysis using GWAS and Allele-Specific Expression (ASE). This led to the development of *nf-pegASE* (INRAE, Table 1), a pipeline for ASE analysis that is based on the nf-core template which is intended to be contributed to nf-core. Some pipelines were also developed outside of the nf-core umbrella, like TAGADA^35^ for the annotation and quantification of novel long non-coding and coding RNA transcripts and *BovReg/nf-cage*^36^ for transcript annotation. Thanks to the standardisation provided by Nextflow, these non-nf-core pipelines can nonetheless provide new modules and elements lending themselves to nf-core integration. Overall, most pipeline methods were widely shared across the various consortium members and while nf-core dominates, other solutions were also part of a very active exchange process (Fig. 2b, Supplementary Table 3) with 4 of the 17 pipelines being used by at least three EuroFAANG consortiums.

### Conclusion and perspective

The rapid growth of the nf-core community demonstrates the importance of strict best practice guidelines bundled with tooling for developers and users. The module and subworkflow infrastructure facilitates collaborative development both within nf-core and between external pipelines, leading to improved code quality and long-term maintenance. nf-core standards and community support enable faster development, improved interoperability, standardisation, and collaboration. As adoption spreads from traditional bioinformatics to other scientific fields these standards continue to enable other distributed communities to work together effectively and produce more reliable research.

In this report, we present the process of adopting a bioinformatics pipeline standard at the level of a group of consortia involving over 80 institutions and companies, with a primary focus on computational analysis. We argue here for soft standards, meaning a framework that accommodates a continuum between the current state of an implementation and the most desirable one. We found nf-core to be a good trade-off as it is based on Nextflow, which can wrap any existing code regardless of its implementation language. Strict adherence to nf-core best-practices is not mandatory for a Nextflow pipeline to be functional, allowing for progressive development towards nf-core standards. The extra development effort is offset by secondary benefits resulting from community input: nf-core pipelines are more visible, and have more accessible documentation. They benefit from community support, validation, bug fixes, and extension. These factors are especially important when considering long-term sustainability.

The EuroFAANG collaboration with nf-core has been a great success, resulting in mutual benefits and truly FAIR science. To foster more collaborations of this kind, nf-core has started a new initiative of “special interest groups”, where researchers with a common focus can collaborate on standardised usage of nf-core pipelines. We hope that these groups will work orthogonally to existing groups working on specific pipelines, encouraging improved collaboration and standardisation between scientists worldwide.

