## Supplementary Material for "Empowering bioinformatics communities with Nextflow and nf-core"

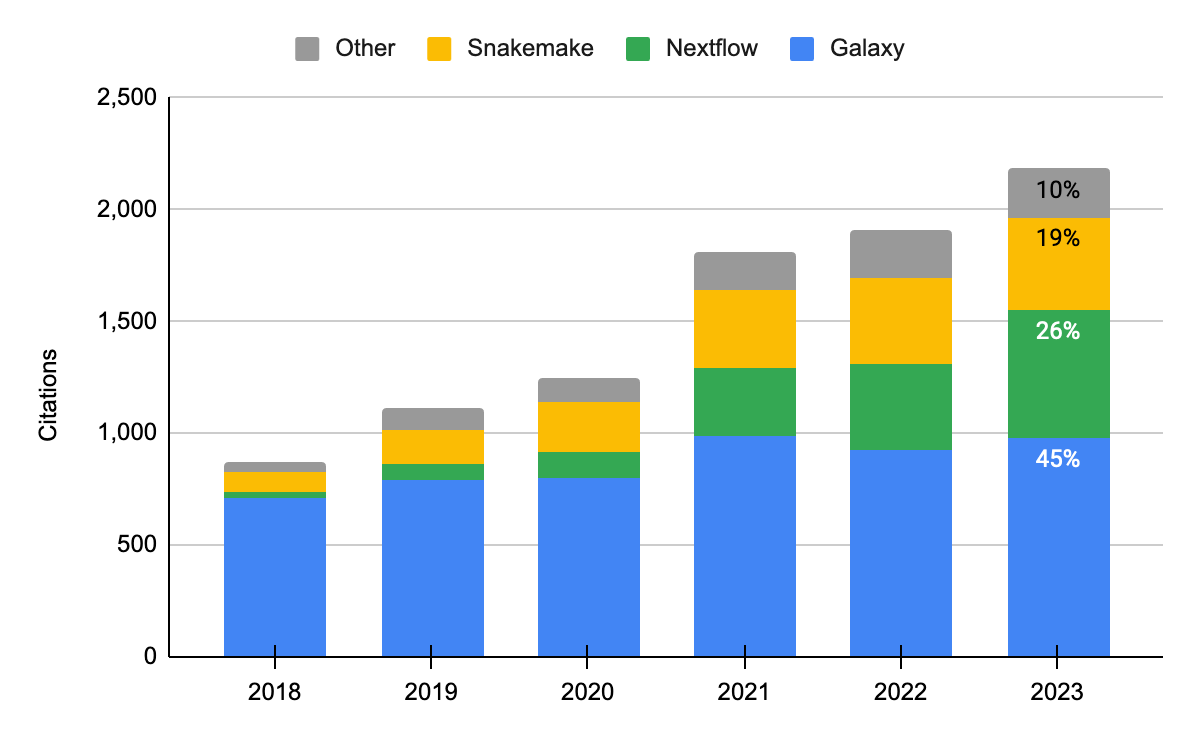

**Supplementary Figure 1: Scopus citation counts for bioinformatics workflow management systems.** Sum of citations of the major publications of Galaxy, Nextflow, and Snakemake between 2018 and 2023 (Data in Supplementary Table 2).

| **WFMS** | **Title** | **2018** | **2019** | **2020** | **2021** | **2022** | **2023** | **Total** |
| --- | --- | --- | --- | --- | --- | --- | --- | --- |
| Galaxy | The Galaxy platform for accessible, reproducible and collaborative biomedical analyses: 2022 update^1^ |  |  |  |  | 10 | 140 | **150** |
| Galaxy | Correction to ‘The Galaxy platform for accessible, reproducible and collaborative biomedical analyses: 2022 update‘ ^2^ | *non-referenced* | | | | | | |
| Galaxy | The Galaxy platform for accessible, reproducible and collaborative biomedical analyses: 2020 update^3^ |  |  | 8 | 95 | 156 | 119 | **378** |
| Galaxy | Corrigendum: The Galaxy platform for accessible, reproducible and collaborative biomedical analyses: 2020 update^4^ | *non-referenced* | | | | | | |
| Galaxy | The Galaxy platform for accessible, reproducible and collaborative biomedical analyses: 2018 update^5^ | 64 | 398 | 616 | 772 | 791 | 703 | **3,344** |
| Galaxy | The Galaxy platform for accessible, reproducible and collaborative biomedical analyses: 2016 update^6^ | 434 | 350 | 262 | 257 | 192 | 143 | **1,638** |
| Galaxy | Galaxy: a comprehensive approach for supporting accessible, reproducible, and transparent computational research in the life sciences^7^ | 310 | 268 | 161 | 167 | 133 | 104 | **1,143** |
| Galaxy | Galaxy: a web‐based genome analysis tool for experimentalists^8^ | 123 | 77 | 56 | 59 | 37 | 30 | **382** |
| Galaxy | Galaxy: a platform for interactive large-scale genome analysis^9^ | 193 | 145 | 136 | 118 | 85 | 67 | **744** |
| KNIME | KNIME for reproducible cross-domain analysis of life science data^10^ | 17 | 25 | 30 | 34 | 25 | 32 | **163** |
| KNIME | KNIME-the Konstanz information miner: version 2.0 and beyond^11^ | 279 | 276 | 306 | 312 | 316 | 274 | **1,763** |
| KNIME | Data analysis, machine learning and applications^12^ | 14 | 18 | 27 | 15 | 20 | 12 | **106** |
| Nextflow | The nf-core framework for community-curated bioinformatics pipelines^13^ |  | 7 | 50 | 159 | 273 | 430 | **919** |
| Nextflow | Nextflow enables reproducible computational workflows^14^ | 71 | 144 | 195 | 307 | 370 | 752 | **1,839** |
| Snakemake | Sustainable data analysis with Snakemake^15^ |  |  | 2 | 81 | 234 | 398 | **715** |
| Snakemake | Corrigendum: Snakemake—a scalable bioinformatics workflow engine^16^ | *non-referenced* | | | | | | |
| Snakemake | Snakemake—a scalable bioinformatics workflow engine^17^ | 170 | 271 | 392 | 498 | 436 | 392 | **2,159** |
| Bpipe | Bpipe: a tool for running and managing bioinformatics pipelines^18^ | 24 | 25 | 22 | 19 | 7 | 10 | **107** |
| Pachyderm | Container-based bioinformatics with Pachyderm^19^ | 3 | 17 | 9 | 12 | 5 | 11 | **57** |
| SciPipe | SciPipe: a workflow library for agile development of complex and dynamic bioinformatics pipelines^20^ | 1 | 5 | 7 | 12 | 7 | 6 | **38** |
| Cromwell | Full-stack genomics pipelining with GATK4 + WDL + Cromwell^21^ | 6 | 13 | 9 | 24 | 20 | 14 | **86** |
| Toil (CWL) | Toil enables reproducible, open source, big biomedical data analyses^22^ | 40 | 100 | 95 | 152 | 251 | 250 | **888** |
| CWL | Common Workflow Language, v1. 0^23^ | 21 | 36 | 36 | 34 | 22 | 15 | **164** |
| **Total** |  | **1,770** | **2,175** | **2,419** | **3,127** | **3,390** | **3,902** | **16,783** |
| Σ Galaxy |  | 1,124 | 1,238 | 1,239 | 1,468 | 1,404 | 1,306 | **7,779** |
| Σ Nextflow |  | 71 | 151 | 245 | 466 | 643 | 1,182 | **2,758** |
| Σ Snakemake |  | 170 | 271 | 394 | 579 | 670 | 790 | **2,874** |

**Supplementary Table 1: Google Scholar citation counts for bioinformatics workflow management systems.** WfMSs were selected based on a recent review^24^. Publications dedicated to these resources were then collected, excluding domain-specific work, and searched on Google Scholar (2024-04-19). The list of citations provided by Google Scholar was then ventilated by year using the range option of Google interface. The corrigendum and erratum are not referenced on Google Scholar.

| **WFMS** | **Title** | **2018** | **2019** | **2020** | **2021** | **2022** | **2023** | **Total** |
| --- | --- | --- | --- | --- | --- | --- | --- | --- |
| Galaxy | The Galaxy platform for accessible, reproducible and collaborative biomedical analyses: 2022 update^1^ |  |  |  |  | 12 | 165 | **177** |
| Galaxy | Correction to ‘The Galaxy platform for accessible, reproducible and collaborative biomedical analyses: 2022 update‘ ^2^ |  |  |  |  |  | 3 | **3** |
| Galaxy | The Galaxy platform for accessible, reproducible and collaborative biomedical analyses: 2020 update^3^ |  |  | 2 | 64 | 94 | 75 | **235** |
| Galaxy | Corrigendum: The Galaxy platform for accessible, reproducible and collaborative biomedical analyses: 2020 update^4^ |  |  | 1 | 6 | 7 | 9 | **23** |
| Galaxy | The Galaxy platform for accessible, reproducible and collaborative biomedical analyses: 2018 update^5^ | 25 | 248 | 398 | 543 | 519 | 482 | **2,215** |
| Galaxy | The Galaxy platform for accessible, reproducible and collaborative biomedical analyses: 2016 update^6^ | 292 | 233 | 174 | 171 | 134 | 100 | **1,104** |
| Galaxy | Galaxy: a comprehensive approach for supporting accessible, reproducible, and transparent computational research in the life sciences^7^ | 204 | 180 | 103 | 106 | 87 | 75 | **755** |
| Galaxy | Galaxy: a web‐based genome analysis tool for experimentalists^8^ | 69 | 48 | 43 | 31 | 26 | 24 | **241** |
| Galaxy | Galaxy: a platform for interactive large-scale genome analysis^9^ | 117 | 83 | 77 | 67 | 50 | 47 | **441** |
| KNIME | KNIME for reproducible cross-domain analysis of life science data^10^ | 12 | 12 | 19 | 29 | 18 | 22 | **112** |
| KNIME | KNIME-the Konstanz information miner: version 2.0 and beyond^11^ | *non-referenced* | | | | | | |
| KNIME | Data analysis, machine learning and applications^12^ | *non-referenced* | | | | | | |
| Nextflow | The nf-core framework for community-curated bioinformatics pipelines^13^ |  |  | 15 | 107 | 163 | 251 | **536** |
| Nextflow | Nextflow enables reproducible computational workflows^14^ | 33 | 68 | 102 | 194 | 214 | 318 | **929** |
| Snakemake | Sustainable data analysis with Snakemake^15^ |  |  | 0 | 39 | 116 | 208 | **363** |
| Snakemake | Corrigendum: Snakemake—a scalable bioinformatics workflow engine^16^ | 1 | 10 | 28 | 44 | 33 | 23 | **139** |
| Snakemake | Snakemake—a scalable bioinformatics workflow engine^17^ | 86 | 148 | 199 | 271 | 236 | 187 | **1,127** |
| Bpipe | Bpipe: a tool for running and managing bioinformatics pipelines^18^ | 10 | 16 | 12 | 12 | 3 | 6 | **59** |
| Pachyderm | Container-based bioinformatics with Pachyderm^19^ | 0 | 8 | 6 | 10 | 4 | 4 | **32** |
| SciPipe | SciPipe: a workflow library for agile development of complex and dynamic bioinformatics pipelines^20^ | 0 | 1 | 5 | 8 | 1 | 2 | **17** |
| Cromwell | Full-stack genomics pipelining with GATK4 + WDL + Cromwell^21^ | *non-referenced* | | | | | | |
| Toil (CWL) | Toil enables reproducible, open source, big biomedical data analyses^22^ | 24 | 62 | 59 | 111 | 190 | 182 | **628** |
| CWL | Common Workflow Language, v1. 0^23^ | *non-referenced* | | | | | | |
| **Total** |  | **873** | **1,117** | **1,243** | **1,813** | **1,907** | **2,183** | **9,136** |
| Σ Galaxy |  | 707 | 792 | 798 | 988 | 929 | 980 | **5,194** |
| Σ Nextflow |  | 33 | 68 | 117 | 301 | 377 | 569 | **1,465** |
| Σ Snakemake |  | 87 | 158 | 227 | 354 | 385 | 418 | **1,629** |

**Supplementary Table 2: Scopus Scholar citation counts for bioinformatics workflow management systems.** WfMSs were selected based on a recent review^24^. Publications dedicated to these resources were then collected, excluding domain-specific work, and searched on Scopus (2024-04-19). The reported counts are the ones provided by Scopus in the annual breakdown section. Publications indicated as non-referenced were not found in Scopus.

| Institution | Consortium | Analysis | Language/ Standard | Github / DOI |
| --- | --- | --- | --- | --- |
| HCMR | AQUA-FAANG | RNA-Seq: Embryonic development in gilthead seabream - transcriptome | nf-core | nf-core/rnaseq |
| HCMR | AQUA-FAANG | ATAC-Seq: Embryonic development in gilthead seabream - chromatin accessibility dynamics | nf-core | nf-core/atacseq |
| HCMR | AQUA-FAANG | RNA-Seq: Transcriptome dynamics after immune stimulation in gilthead seabream | nf-core | nf-core/rnaseq |
| HCMR | AQUA-FAANG | ATAC-Seq: Chromatin accessibility dynamics after immune stimulation in gilthead seabream | nf-core | nf-core/atacseq |
| INRAE | AQUA-FAANG | RNA-Seq: Transcriptome dynamics after immune stimulation or experimental infection in rainbow trout | nf-core | nf-core/rnaseq |
| INRAE | AQUA-FAANG | ATAC-Seq: Chromatin accessibility dynamics after immune stimulation or experimental infection in rainbow trout | nf-core | nf-core/atacseq |
| NMBU | AQUA-FAANG | RNA-Seq: Panel of tissues in Atlantic salmon and rainbow trout - transcriptome | nf-core | nf-core/rnaseq |
| NMBU | AQUA-FAANG | ATAC-Seq: Panel of tissues in Atlantic salmon and rainbow trout - chromatin accessibility dynamics | nf-core | nf-core/atacseq |
| NMBU | AQUA-FAANG | ChIP-Seq: Panel of tissues in Atlantic salmon and rainbow trout - histone mark dynamics | nf-core | nf-core/chipseq |
| NMBU | AQUA-FAANG | RNA-Seq: Panel of tissues in Atlantic salmon and rainbow trout - transcriptome | nf-core | nf-core/rnaseq |
| NMBU | AQUA-FAANG | ATAC-Seq: Panel of tissues in Atlantic salmon and rainbow trout - chromatin accessibility dynamics | nf-core | nf-core/atacseq |
| NMBU | AQUA-FAANG | ChIP-Seq: Panel of tissues in Atlantic salmon and rainbow trout - histone mark dynamics | nf-core | nf-core/chipseq |
| PAN | AQUA-FAANG | RNA-Seq: Panel of tissues in common carp - transcriptome | nf-core | nf-core/rnaseq |
| PAN | AQUA-FAANG | ATAC-Seq: Panel of tissues in common carp - chromatin accessibility dynamics | nf-core | nf-core/atacseq |
| PAN | AQUA-FAANG | ChIP-Seq: Panel of tissues in common carp - histone mark dynamics | nf-core | nf-core/chipseq |
| UEDIN | AQUA-FAANG | RNA-Seq: Embryonic development in Atlantic salmon and rainbow trout - transcriptome | nf-core | nf-core/rnaseq |
| UEDIN | AQUA-FAANG | ATAC-Seq: Embryonic development in Atlantic salmon and rainbow trout - chromatin accessibility dynamics | nf-core | nf-core/atacseq |
| UEDIN | AQUA-FAANG | ChIP-Seq: Embryonic development in Atlantic salmon and rainbow trout - histone mark dynamics | nf-core | nf-core/chipseq |
| UNIABDN | AQUA-FAANG | RNA-Seq: Transcriptome dynamics after immune stimulation in Atlantic salmon | nf-core | nf-core/rnaseq |
| UNIABDN | AQUA-FAANG | ATAC-Seq: Chromatin accessibility dynamics after immune stimulation in Atlantic salmon | nf-core | nf-core/atacseq |
| UNIABDN | AQUA-FAANG | ChIP-Seq: Histone mark dynamics after immune stimulation in Atlantic salmon | nf-core | nf-core/chipseq |
| UNIPD | AQUA-FAANG | RNA-Seq: Embryonic development in European seabass - transcriptome | nf-core | nf-core/rnaseq |
| UNIPD | AQUA-FAANG | ATAC-Seq: Embryonic development in European seabass - chromatin accessibility dynamics | nf-core | nf-core/atacseq |
| UNIPD | AQUA-FAANG | ChIP-Seq: Embryonic development in European seabass - histone mark dynamics | nf-core | nf-core/chipseq |
| UNIPD | AQUA-FAANG | RNA-Seq: Panel of tissues in European seabass - transcriptome | nf-core | nf-core/rnaseq |
| UNIPD | AQUA-FAANG | ATAC-Seq: Panel of tissues in European seabass - chromatin accessibility dynamics | nf-core | nf-core/atacseq |
| UNIPD | AQUA-FAANG | ChIP-Seq: Panel of tissues in European seabass - histone mark dynamics | nf-core | nf-core/chipseq |
| UNIPD | AQUA-FAANG | RNA-Seq: Transcriptome dynamics after immune stimulation in European seabass | nf-core | nf-core/rnaseq |
| UNIPD | AQUA-FAANG | ATAC-Seq: Chromatin accessibility dynamics after immune stimulation in European seabass | nf-core | nf-core/atacseq |
| UNIPD | AQUA-FAANG | ChIP-Seq: Histone mark dynamics after immune stimulation in European seabass | nf-core | nf-core/chipseq |
| UoB | AQUA-FAANG | ATAC-Seq: Embryonic development in common carp - chromatin accessibility dynamics | nf-core | nf-core/atacseq |
| UoB | AQUA-FAANG | ChIP-Seq: Embryonic development in common carp - histone mark dynamics | nf-core | nf-core/chipseq |
| USC | AQUA-FAANG | RNA-Seq: Embryonic development in turbot - transcriptome | nf-core | nf-core/rnaseq |
| USC | AQUA-FAANG | ATAC-Seq: Embryonic development in turbot - chromatin accessibility dynamics | nf-core | nf-core/atacseq |
| USC | AQUA-FAANG | ChIP-Seq: Embryonic development in turbot - histone mark dynamics | nf-core | nf-core/chipseq |
| USC | AQUA-FAANG | RNA-Seq: Panel of tissues in turbot - transcriptome | nf-core | nf-core/rnaseq |
| USC | AQUA-FAANG | ATAC-Seq: Panel of tissues turbot - chromatin accessibility dynamics | nf-core | nf-core/atacseq |
| USC | AQUA-FAANG | ChIP-Seq: Panel of tissues in turbot - histone mark dynamics | nf-core | nf-core/chipseq |
| USC | AQUA-FAANG | RNA-Seq: Transcriptome dynamics after immune stimulation in turbot | nf-core | nf-core/rnaseq |
| USC | AQUA-FAANG | ATAC-Seq: Chromatin accessibility dynamics after immune stimulation in turbot | nf-core | nf-core/atacseq |
| USC | AQUA-FAANG | ChIP-Seq: Histone mark dynamics after immune stimulation in turbot | nf-core | nf-core/chipseq |
| FBN | BovReg | ChIRP | bash |  |
| FBN | BovReg | ChIRP | nf-core | nf-core/chipseq |
| FBN | BovReg | circRNAs | bash |  |
| FBN | BovReg | eQTLs, sQTLs | Nextflow | BovReg/BovReg_eQTL |
| FBN | BovReg | Post-GWAS | Bash |  |
| FBN | BovReg | RRBS | nf-core | nf-core/methylseq |
| FBN | BovReg | RNA-Seq | nf-core | nf-core/rnaseq |
| FMV | BovReg | miRNAs expression | Perl | andreiaamaral/ IsomiR-Window |
| GIGA | BovReg | ATAC-Seq: Open chromatin region identification and quantification | nf-core | nf-core/atacseq |
| GIGA | BovReg | Protein binding region identification and quantification | nf-core | nf-core/chipseq |
| GIGA | BovReg | miRNAs expression | nf-core | nf-core/smrnaseq |
| GIGA | BovReg | Transcript annotation and quantification (mRNA and lncRNA) | nf-core | BovReg/rnaseq |
| GIGA | BovReg | RNA-Seq: Transcript annotation and quantification (long-read cDNA sequencing / Oxford Nanopore technology) | nf-core | nf-core/nanoseq |
| INRAE | BovReg | Hi-C: Pairs of proximal DNA region identification and quantification | nf-core | nf-core/hic |
| INRAE | BovReg | SNPs/indels/SVs: Variant detection in 5 bovine cell lines | nf-core | nf-core/sarek |
| INRAE | BovReg | EM-Seq: Analysis of Enzymatic Methylation sequencing data for five bovine cell lines | nf-core | nf-core/methylseq |
| LUKE | BovReg | GWAS | Snakemake |  |
| LUKE | BovReg | RNA-Seq | Snakemake | [10.5281/zenodo.7743750](https://doi.org/10.5281/zenodo.7743750) |
| LUKE | BovReg | miRNA-Seq | Snakemake | [10.5281/zenodo.7707383](https://doi.org/10.5281/zenodo.7707383) |
| UEDIN | BovReg | CAGE-seq | nf-core | BovReg/nf-cage |
| UEDIN | BovReg | SNP heritability: Heritability partitioning by LD-score regression | Nextflow |  |
| UEDIN | BovReg | TWAS: Genotype imputation and gene expression prediction model generation | Nextflow |  |
| INRAE-INSERM | GENE-SWitCH | ATAC-Seq: Open chromatin region identification and quantification | nf-core | nf-core/atacseq |
| INRAE-INSERM | GENE-SWitCH | Hi-C: Pairs of proximal DNA region identification and quantification | nf-core | nf-core/hic |
| INRAE-INSERM | GENE-SWitCH | RNA-Seq: Transcript and gene annotation and expression | Nextflow | FAANG/analysis-TAGADA |
| INRAE-INSERM | GENE-SWitCH | Small RNA-Seq: Small non-coding transcript annotation and expression | nf-core | nf-core/smrnaseq |
| UEDIN | GENE-SWitCH | Iso-Seq: Transcript and gene annotation | nf-core | nf-core/isoseq |
| WUR | GENE-SWitCH | ChIP-seq: Protein binding region identification and quantification | nf-core | nf-core/chipseq |
| WUR | GENE-SWitCH | WGBS: Methylated region identification and quantification | nf-core | FAANG/GSM-pipeline |
| INRAE-Institut Agro | GEroNIMO | eQTLs: TensorQTL | bash |  |
| INRAE-Institut Agro | GEroNIMO | EM-Seq / WGBS / RRBS: DNA methylation analyses (Enzymatic Methyl-Seq and Bisulfite sequencing) | nf-core | nf-core/methylseq |
| INRAE-Institut Agro | GEroNIMO | Small RNA-Seq: Small non-coding transcript annotation and expression | nf-core | nf-core/smrnaseq |
| INRAE-Institut Agro | GEroNIMO | RNA-Seq: Transcript and gene annotation and expression | nf-core | nf-core/rnaseq |
| INRAE-Institut Agro | GEroNIMO | RNA-Seq: variation identifier from RNA-Seq | nf-core | nf-core/rnavar |
| INRAE-Institut Agro | GEroNIMO | Allele specific expression: Measurement of haplotypic expression in RNAseq assays | nf-core | cguyomar/nf-pegASE |
| IRTA | HoloRuminant | Metagenome: Recovering public metataxonomic and Shotgun Metagenomic datasets | nf-core | nf-core/fetchngs |
| GIGA | RUMIGEN | SNPs/indels/SVs: Variant detection. Accessing the impact of genome-editing and on genome integrity during early embryonic developmental stages | nf-core | nf-core/sarek |

**Supplementary Table 3: Examples of analyses conducted in EuroFAANG projects.** Non-exhaustive list of analyses performed by different groups in the EuroFAANG consortia together with the utilised analysis workflow. Indicated is the workflow’s implementation language or “nf-core” if a Nextflow pipeline is built with the nf-core template.

| **Event** | | **Views** | | | | | | | **Total** |
| --- | --- | --- | --- | --- | --- | --- | --- | --- | --- |
| **October 2022 training** |  | Asia-Pacific | | Europe, Mid. East, Africa | | | America | |  |
|  | Day 1 | 1,246 | | 1,424 | | | 1,138 | | 3,808 |
|  | Day 2 | 301 | | 682 | | | 614 | | 1,597 |
|  | Day 3 | 283 | | 521 | | | 552 | | 1,356 |
| **March 2023 training** |  | English | Portuguese | | Spanish | French | | Hindi |  |
|  | Session 1 | 7,316 | 584 | | 663 | 642 | | 596 | 9,801 |
|  | Session 2 | 2,388 | 149 | | 254 | 202 | | 170 | 3,163 |
|  | Session 3 | 1,635 | 125 | | 169 | 181 | | 80 | 2,190 |
|  | Session 4 | 887 | 106 | | 129 | 104 | | 69 | 1,295 |
| **September 2023 training** |  | English | | | | | | |  |
|  | Foundational session 1 | 6,618 | | | | | | | 6,618 |
|  | Foundational session 2 | 1,427 | | | | | | | 1,427 |
|  | Foundational session 3 | 712 | | | | | | | 712 |
|  | Hands-on training | 1,692 | | | | | | | 1,692 |
|  | Advanced training 1 | 1,136 | | | | | | | 1,136 |
|  | Advanced training 2 | 364 | | | | | | | 364 |
|  |  | **Grand Total** | | | | | | | **35,159** |

**Supplementary Table 4: nf-core outreach over the 2022-2023 period.** Recordings of the various sessions were posted on YouTube and the figures indicate the number of views as of 2024-02-15).
